# ZZZ3 protects human embryonic stem cells from nucleolar stress by boosting mTOR/ribosome pathway

**DOI:** 10.1101/2023.09.08.556837

**Authors:** Michela Lo Conte, Valeria Lucchino, Stefania Scalise, Clara Zannino, Maria Stella Murfuni, Chiara Cicconetti, Luana Scaramuzzino, Danilo Swann Matassa, Anna Procopio, Giovanni Cuda, Elvira Immacolata Parrotta

**Author notes:** **Corresponding author** (Prof. Giovanni Cuda). Equal contributors (Michela Lo Conte, Valeria Lucchino).

## Abstract

Embryonic stem cells (ESCs) are defined as stem cells with self-renewing and differentiation capabilities. These unique properties are tightly regulated and controlled by complex genetic and molecular mechanisms whose understanding is essential for both basic and translational research. A large number of studies have mostly focused on understanding the molecular mechanisms governing pluripotency and differentiation of ESCs, while the regulation of proliferation has received comparably less attention. In mouse ESCs, pluripotency and proliferation can be independent processes meaning that it is possible for mouse ESCs to maintain their pluripotent state without actively proliferating. Here, we investigate the role of ZZZ3 (Zinc Finger ZZ-Type Containing 3) function in human ESCs homeostasis. We found that knockdown of ZZZ3 strongly decreases ribosome biogenesis, translation, and mTOR signaling leading to nucleolar stress and significant reduction of cell proliferation. This process occurs without affecting pluripotency, suggesting that ZZZ3-depleted ESCs enter a dormant-like state and that proliferation and pluripotency can be uncoupled also in human ESCs.

## Introduction

Embryonic stem cells (ESCs) are pluripotent stem cells derived from the inner cell mass of a developing embryo at the blastocyst stage (1,2). These cells have the unique ability to self-renew indefinitely while maintaining their pluripotency and the remarkable ability to differentiate into the three primary germ layers (ectoderm, mesoderm, and endoderm) (3,4). This dynamic nature is due to the ability of ESCs to undergo large fluctuations in gene expression, resulting in a variability of the protein landscape (5). A comprehensive understanding of the molecular and cellular mechanisms involved in the regulation of pluripotency is important for both basic and translational research. To date, a large body of literature has mostly focused on understanding the transcriptional regulatory networks governing pluripotency and differentiation of ESCs (6–9), while understanding of how proliferation is regulated and controlled in these cells is still limited. Research in mouse ESCs has indicated that pluripotency and proliferation can be regulated independently and the mTOR pathway seems to be a key player in governing developmental pausing in mouse ESCs. Paused blastocysts display a significant reduction of mTOR activity, gene expression, cell growth and proliferation, and translation while keeping their pluripotency intact (10). Genetic inactivation of Raptor, an mTORC1-specific protein, was demonstrated to be sufficient to reduce proliferation of mESCs without affecting pluripotency (11). Embryonic development arrest at the blastocyst stage induces the embryo to enter into the diapause, a reversible state of biosynthetic dormancy where the Wnt/β-catenin and Esrrb play a crucial role (12,13). Cell growth and proliferation are the results of complex biosynthetic and metabolic processes which, among others, require proper ribosome biogenesis and translation. ZZZ3 is a histone H3 reader and one of the largest subunits within the ATAC (Ada-two-A-containing) complex. Previous work showed that ZZZ3 regulates ribosomal gene expression in mouse ESCs and in human adenocarcinoma cell lines (14,15). Here, we aim at investigating the ATAC-independent function of ZZZ3 in regulating ribosomal genes in human pluripotent stem cells. We used a combination of interactome, transcriptome, and cellular analyses to show that ZZZ3 protects hESCs from nucleolar stress by acting on mTOR/ribosome biogenesis pathway. Interestingly, the knockdown of ZZZ3 resulted in a significant reduction of hESCs proliferation while pluripotency and differentiation were preserved, suggesting that a pluripotent state of biosynthetic quiescence may occur *in vitro* in human ESCs.

## Results

### ZZZ3 interacts with ribosomal proteins and co-localizes in the nucleolus with fibrillarin

To characterize the ZZZ3 biological network in hESCs, we performed a co-immunoprecipitation assay using an anti-ZZZ3 antibody and co-precipitated proteins were identified using nanoscale liquid chromatography coupled to tandem mass spectrometry (nanoLC-MS/MS). We used two human ESC lines, WA17 and RUES, herein referred to as ESC-1 and ESC-2, respectively. Using the Perseus software for statistical analysis, we identified 400 proteins co-purified with ZZZ3 in ESC-1 and 742 proteins co-purified with ZZZ3 in ESC-2. ZZZ3 interactors were selected based on statistically significant differences (*p* value < 0.05 corrected using the Benjamini-Hochberg procedure and FC ≥ 2.5) compared to samples immunoprecipitated with control IgG. GO (Gene Ontology) enrichment analysis revealed that ribosome biogenesis, rRNA processing, rRNA metabolism, RNP complex biogenesis, RNP complex assembly, and cytoplasmic translation were the most overrepresented annotations for BP (Biological Processes) (Fig. 1A; Table S1). The Small Subunit (SSU) Processome, spliceosome complex, and ribosome were the most dominant terms in the Cellular Components (CC) GO hierarchy, while the Molecular Function (MF) category was enriched by terms such as structural constituent of ribosome, rRNA binding, snRNA binding, helicase activity (Fig. S1A; Table S1). Protein-protein interactions (PPI) networks analysis of the proteins directly interacting with ZZZ3 (FC > 10) and enriching biological processes with higher scores include ribosomal protein components of the 40S (RPS8, RPS15, RPS15A) and 60S subunits (RPL27, RPL27A, RPL36) (Fig. 1B). The major biological processes enriched by ZZZ3 interacting proteins are illustrated in the PPI network (Fig. 1C). Since ribosome biogenesis occurs primarily in the cell’s nucleolus, we used immunostaining to mark the co-expression of ZZZ3 and FBL, a nucleolar protein actively involved in pre-rRNA processing, pre-rRNA methylation, and ribosome assembly (24,25), in wild-type ESC-1 and −2. The intensity-based correlation analysis conducted with PCC and MOC (Pearson’s and Manders’ correlation coefficients, respectively) analysis showed that ZZZ3 and FBL strongly co-localize in the nucleolus of hESCs (Fig. S1B). In line with previous studies linking ZZZ3 to the regulation of ribosomal gene expression (14,15), we found that many interactors of ZZZ3 are ribosomal proteins and that ZZZ3 co-localized with FBL in the nucleolus.

**Figure 1.**
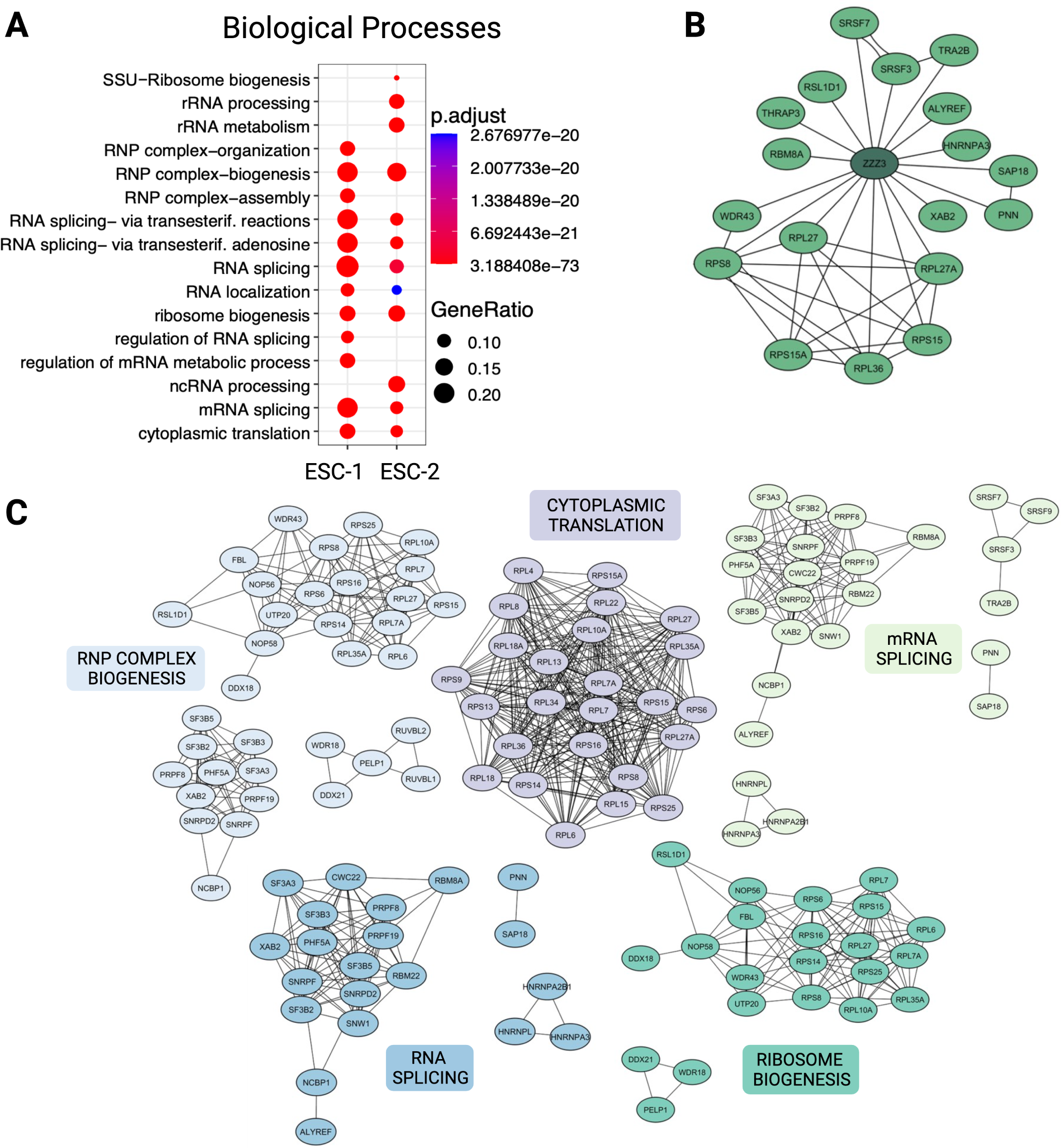
ZZZ3 interacts with proteins involved in cytoplasmic translation, ribosome biogenesis, RNP complex biogenesis, and RNA processing. **A.** Gene ontology (GO) enrichment analysis for biological processes (BP) conducted on genes selected on the basis of their *p*-values (*p*-value < 0.05 corrected using the Benjamini-Hochberg procedure) and fold-change (FC ≥ 2.5). GO was conducted in *R*, using the Bioconductor package. **B.** Protein–protein interaction (PPI) network of the top differentially expressed genes (DEGs) (FC ≥ 10) enriching the BP with the highest gene ratio in ESC-1 and ESC-2. Edges indicate known interactions between DEGs and ZZZ3. **C.** PPI network of all members of the top five BP with the highest gene ratio in ESC-1 and ESC-2. All PPI networks were constructed using the Cytoscape software (version 3.10.0).

### Pluripotency and differentiation are not compromised upon ZZZ3 deficiency

Pluripotent ESCs are highly proliferative, meaning that the ribosome biogenesis must occur at a high rate to sustain their growth and proliferation. If ZZZ3 interacts with and may be associated with the regulation of genes encoding ribosomal protein, what are the effects of ZZZ3 depletion on pluripotency? To answer this question, we stably knocked down the ZZZ3 expression in ESC-1 and ESC-2 lines using piggyBac (PB) vectors (Fig. S2A and S2B). Three independent PB-shRNAs were used to achieve ZZZ3 silencing; shRNA#2 was selected for further experiments based on its high KD efficiency in both ESC lines (Fig. S2C and S2D). First, we analyzed the expression of alkaline phosphatase (AP) to assess the proportion of undifferentiated cells between ZZZ3 KD and SCR control ESCs. We did not detect differences neither in the AP activity nor in cell morphology between ZZZ3 KD ESCs and control cells (Fig. 2A). Pluripotency is stabilized by a triad of interconnected pluripotency transcription factors, namely OCT4, SOX2, and NANOG, that cooperatively regulate gene expression (26,27). Quantitative immunoblot and immunostaining confirmed maintenance of OCT4, NANOG and SOX2 expression at the protein level in ZZZ3 KD ESCs (Fig. 2B and 2C). Gene expression analysis by RNA-seq of a set of pluripotency regulator genes including *OCT4, SOX2, NONOG, MYC, LIN28A, LIN28B, BCOR, FOXO1, DPPA2, DPPA4, UTF1, KLF7* revealed a similar expression pattern in modified and control ESCs (Fig. 2D). *OCT4*, *NANOG*, and *SOX2* mRNA expression levels were validated by qRT-PCR (Fig. S2E). To investigate further in this direction, as additional means for pluripotency we analyzed the capacity of ZZZ3 KD hESCs to form Embryoid Bodies (EBs) and differentiate properly. EBs at day 28 of differentiation were immunoassayed with antibodies against Actin, Sox17, and Nestin, specific for mesoderm, endoderm, and ectoderm, respectively. This assay allowed us to conclude that ZZZ3 KD hESCs can differentiate in a fashion similar to control cells providing additional confirmation that ZZZ3 depletion does not impair pluripotency (Fig. S2F).

**Figure 2.**
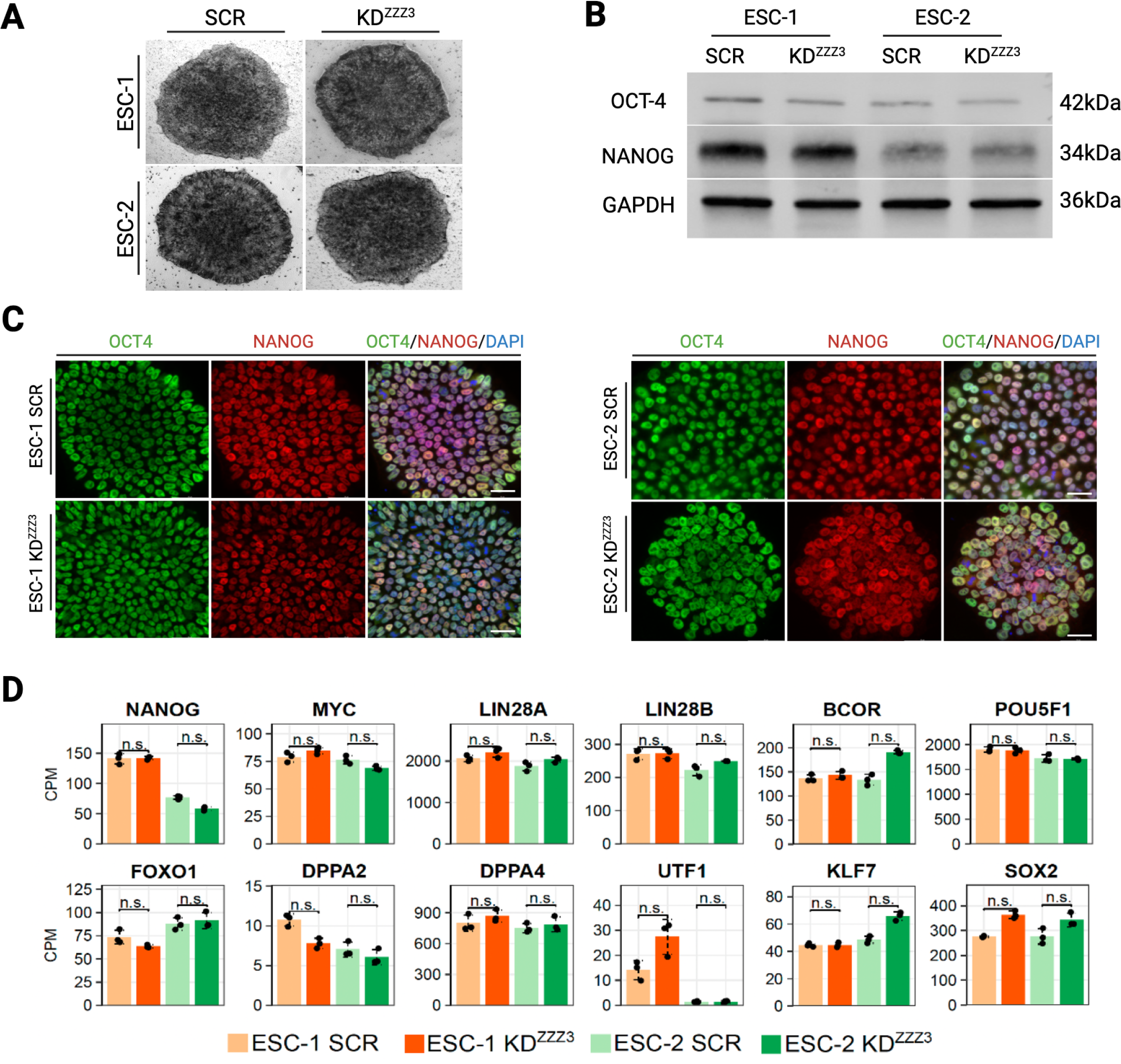
Pluripotency and differentiation are kept intact upon ZZZ3 knockdown. **A.** Alkaline phosphatase (AP) staining of SCR control and ZZZ3 KD ESCs. **B.** Quantitative immunoblot analysis of OCT4 and NANOG expression levels. Full-length blots are shown in Supplementary File S1. **C.** Immunofluorescence analysis of OCT4 and NANOG showing comparable expression of OCT4 and NANOG between ZZZ3-depleted cells and SCR control. Nuclei were counterstained using DAPI. Scale bar, 50 µm. **D**. Barplots of gene expression level from RNA-seq of a set of pluripotency regulators in SCR and ZZZ3 KD ESCs. Bars indicate the mean ± SEM of independent experiments (n = 3) shown as dots. (ANOVA test; n.s.: not significant).

### Knockdown of ZZZ3 induces a dormant-like pluripotent state

Pluripotency and the rate of proliferation are two processes that could be uncoupled. Indeed, during diapause pluripotency is kept intact although proliferation is strongly reduced. Because ZZZ3 interacts with ribosomal proteins and might be involved in the regulation of ribosome biogenesis, which occurs synergistically with cell growth and proliferation, we further wondered what was the effect of its knockdown on ESCs proliferation. Cell proliferation analysis, performed by cell counting at specific time points (24h, 48h, 72h, and 96h) and MTT assay, revealed that ZZZ3 depleted cells were significantly less proliferative compared with control ESCs (Fig. 3A and 3B). These results provide evidence that KD of ZZZ3 negatively affects the capacity of hESCs to undergo cell division. The size of EBs correlates with proliferative capacity of ESCs and cells with faster cell cycle progression are more likely to contribute to larger EBs (28). We measured the diameter of EBs generated from ZZZ3 KD and control hESCs and observed that modified cells generated smaller EBs (Fig. 3C). Next, we performed immunostaining for Ki67, a protein involved in the cell cycle regulation thus serving as a cellular marker for cell proliferation (29,30). We found that the expression of Ki67 is dramatically reduced in ZZZ3 KD hESCs providing further evidence that ZZZ3 is involved in the regulation of human ESCs proliferation (Fig. 3D).

**Figure 3.**
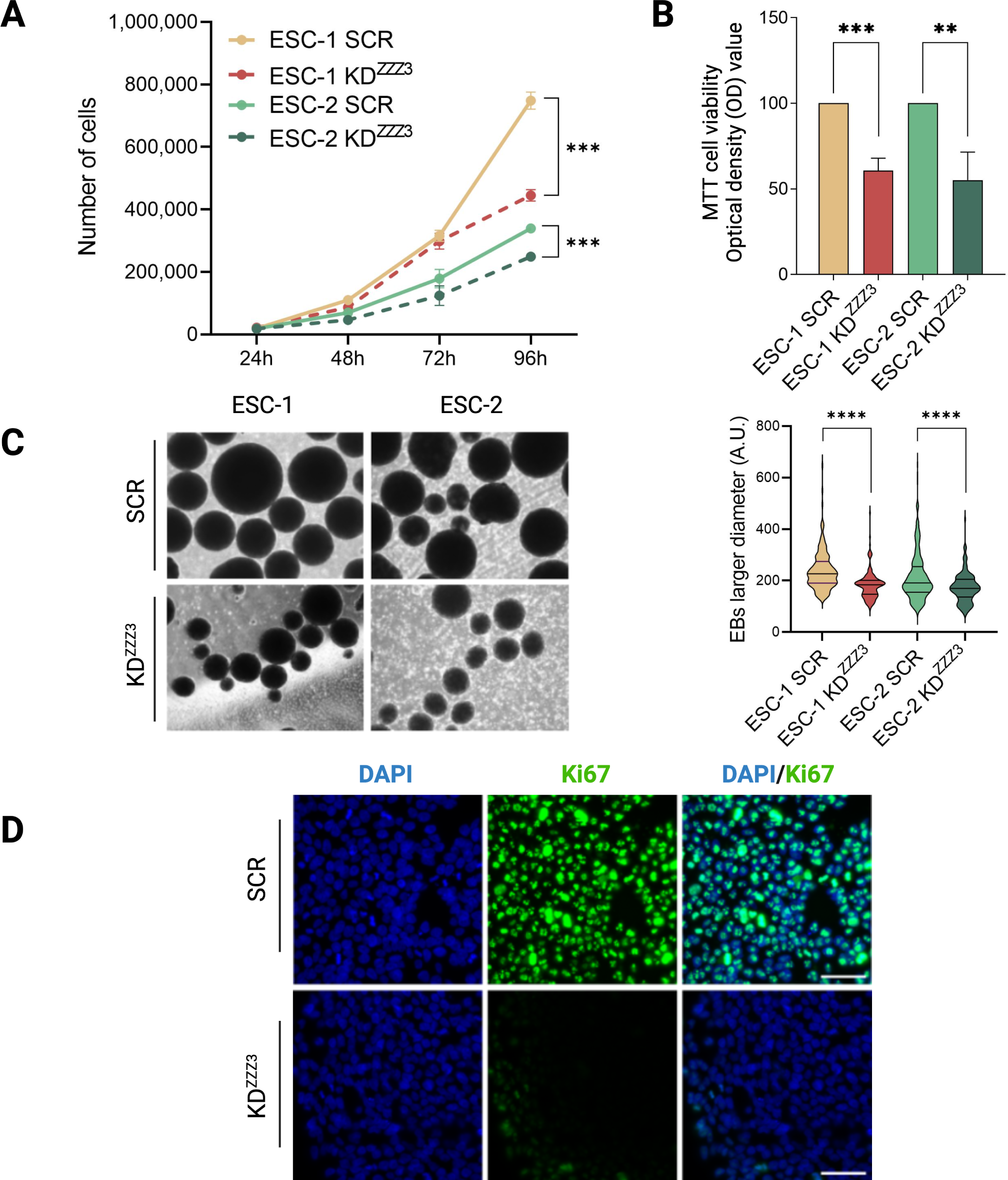
ZZZ3 KD ESCs enter a state of reduced proliferation. **A.** Cell proliferation over a 96-hour time course (n = 3). The initial cell density was 3×10^4^ cells/well. Significance was calculated *vs.* relative SCR ESCs using unpaired two-tailed *t*-test: *** *p* ≤ 0.001. **B.** Optical density (OD) obtained at 570 nm was used for determination of the MTT reduction in ZZZ3 KD ESCs. Data are shown as means ± SEM of three independent experiments. Significance was calculated *vs.* relative SCR ESCs using unpaired two-tailed *t*-test: ** *p* ≤ 0.01, *** *p* ≤ 0.001. **C.** Phase-contrast images of EBs generated from SCR and ZZZ3 KD ESC-1 and ESC-2 (Left panel); Measurement of EBs size showing reduced diameter of EBs generated from ZZZ3 KD ESCs compared to control cells. At least 150 EBs per line were measured. Violin plots indicate the mean ± SEM of three biological replicates; unpaired two-tailed *t*-test was used to calculate the statistical significance *vs*. SCR cells: **** *p* ≤ 0.0001 (Right panel). **D.** Representative images of immunostaining of the proliferation marker Ki67 showing a significant reduction in ZZZ3 KD cells compared to control. Nuclear DNA was stained with DAPI (blue). Scale bar, 50 µm.

### ZZZ3 is associated with ribosome biogenesis and translation

RNA-seq on ZZZ3-KD ESCs was used to gain additional insights into the functional and regulatory roles of ZZZ3 in human ESCs. We identified 275 genes downregulated and 259 genes upregulated upon ZZZ3 knockdown (Fig. 4A; Table S2). Genes with a fold change ≥ 2 and a false discovery rate (FDR) < 0.05 were considered as differentially expressed genes (DEGs) (Fig. 4B). Principal component analysis (PCA) was applied to transcriptome data to investigate the clustering patterns between biological replicates (Fig. 4C). Many of downregulated genes encode small and large ribosomal proteins (RPSs and RPLs) (Fig. 4D). GO enrichment analysis revealed that downregulated genes were predominantly associated with biological processes including post-transcriptional regulation of gene expression, translational initiation, RNA processing, ribosome biogenesis, ribonucleoprotein complex biogenesis, and cytoplasmic translation (Fig. 4E). Taken together, interactome and transcriptome data reveal that ZZZ3 regulates transcription of ribosomal genes but also interacts with ribosomal proteins, suggesting an important role of ZZZ3 in the ribosome biogenesis process.

**Figure 4.**
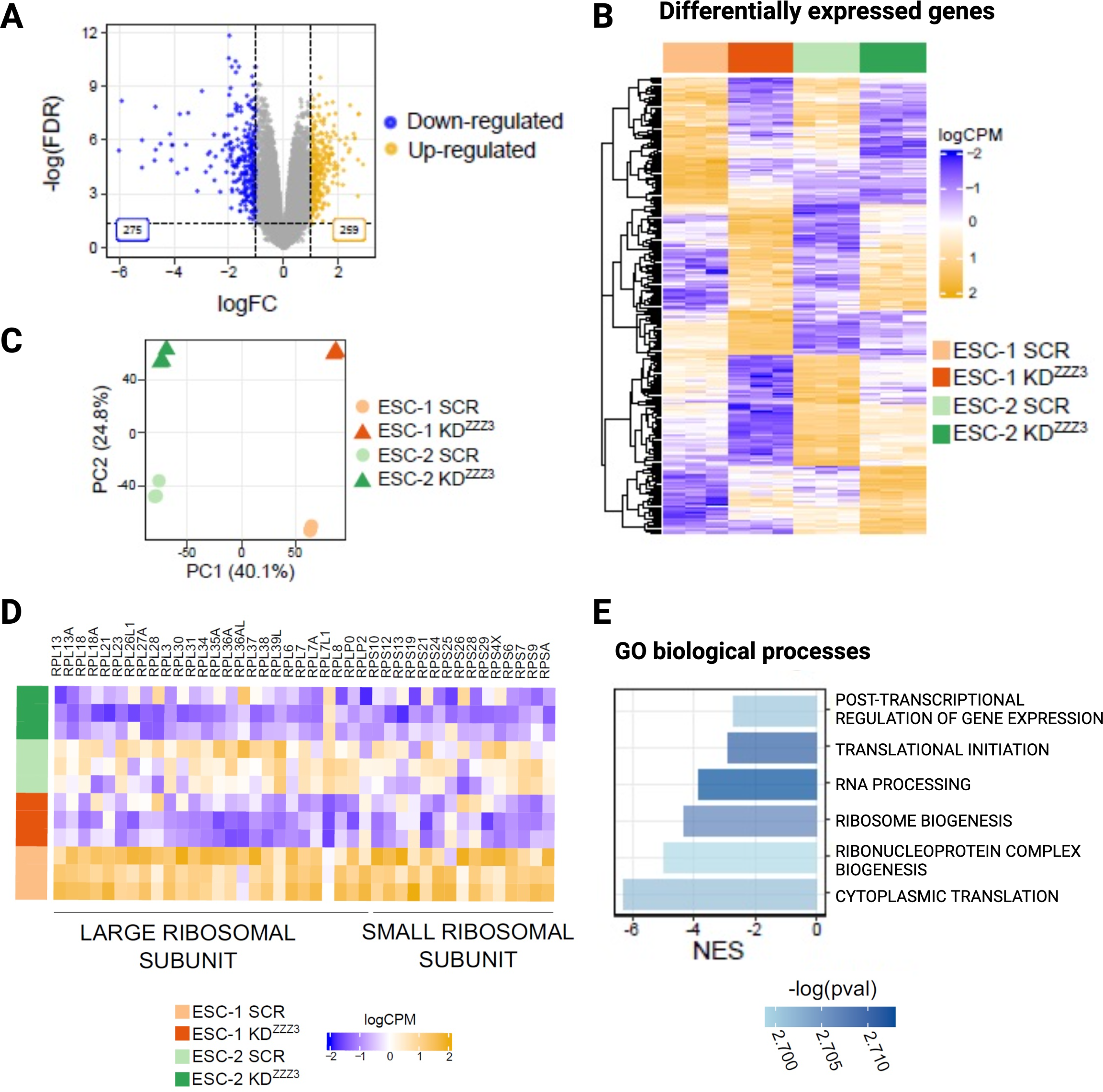
RNA-seq analysis. **A.** Volcano plot depicting differentially expressed genes in ZZZ3 KD ESCs *vs*. SCR control. Blue dots represent downregulated genes (275) while yellow dots represent upregulated genes (259). LogFC and adjusted p-value of the genes are reported. **B**. Heatmap of the DEGs between ZZZ3 KD and SCR condition (Z-score scaled logCPM) by applying the following thresholds: |logFC| > 1 and adjusted p-value < 0.0. Differentially expressed genes are clustered according to a hierarchical clustering procedure based on euclidean distance. **C.** Principal Component Analysis (PCA) of the logCPM gene expression data. **D.** Heatmap of the *Z*-scaled logCPM expression values of genes encoding small and large ribosomal proteins in ZZZ3 KD ESCs and SCR control. **E.** GO Biological Processes gene sets significantly associated with the ZZZ3 KD condition as resulted from the GSEA performed on the logFC*-log(p-value) ranked genes. The Normalized Enriched Score (NES) is reported for each gene set having different color shades according to the −log(p-value).

### ZZZ3 knockdown impairs ribosome biogenesis, translation, and triggers p53 activation

Ribosome biogenesis is a well-orchestrated process involving the coordinated actions of three types of RNA polymerase (RNAP I, RNAP II, and RNAP III) that make up the active ribosomes, machinery of translation (31). To measure and monitor the translation dynamics in ZZZ3 KD ESCs, we used the polysome profiling analysis. This assay clearly showed a reduction in the 80S and polysomes fractions in cells with depletion of ZZZ3 compared to SCR ESCs (Fig. 5A). Western blot analysis for RPL19a and RPS6 was used to validate protein distribution in polysome profiling subfractions (1 to 12) (Fig. S3A). GSEA analysis of transcriptomic data revealed that rRNA processing (NES = - 5.09), translation (NES = - 6.7), and ribosome (NES = - 7.08) are some of the most down regulated biological processes in ZZZ3 KD cells (Fig. 5B and Fig. S3B). Nucleolar stress is a cellular stress mechanism that may result from impaired rRNA synthesis and ribosome biogenesis. Interestingly, we found a more dispersed nucleolar structure in ZZZ3 KD cells compared to SCR ESCs as shown by the immunostaining for FBL, enriched in the dense fibrillar component (DFC) of the nucleolus (Fig. 5C). Impaired ribosome biogenesis can trigger nucleolar stress via a p53-dependent mechanism (32). We thus measured the expression level of p53 and its direct target p21 using immunoblot and immunoassay, respectively (Fig. 5D and 5E). Both were dramatically increased in ZZZ3 KD ESCs, suggesting that nucleolar stress may be associated or concurrent with ZZZ3 depletion.

**Figure 5.**
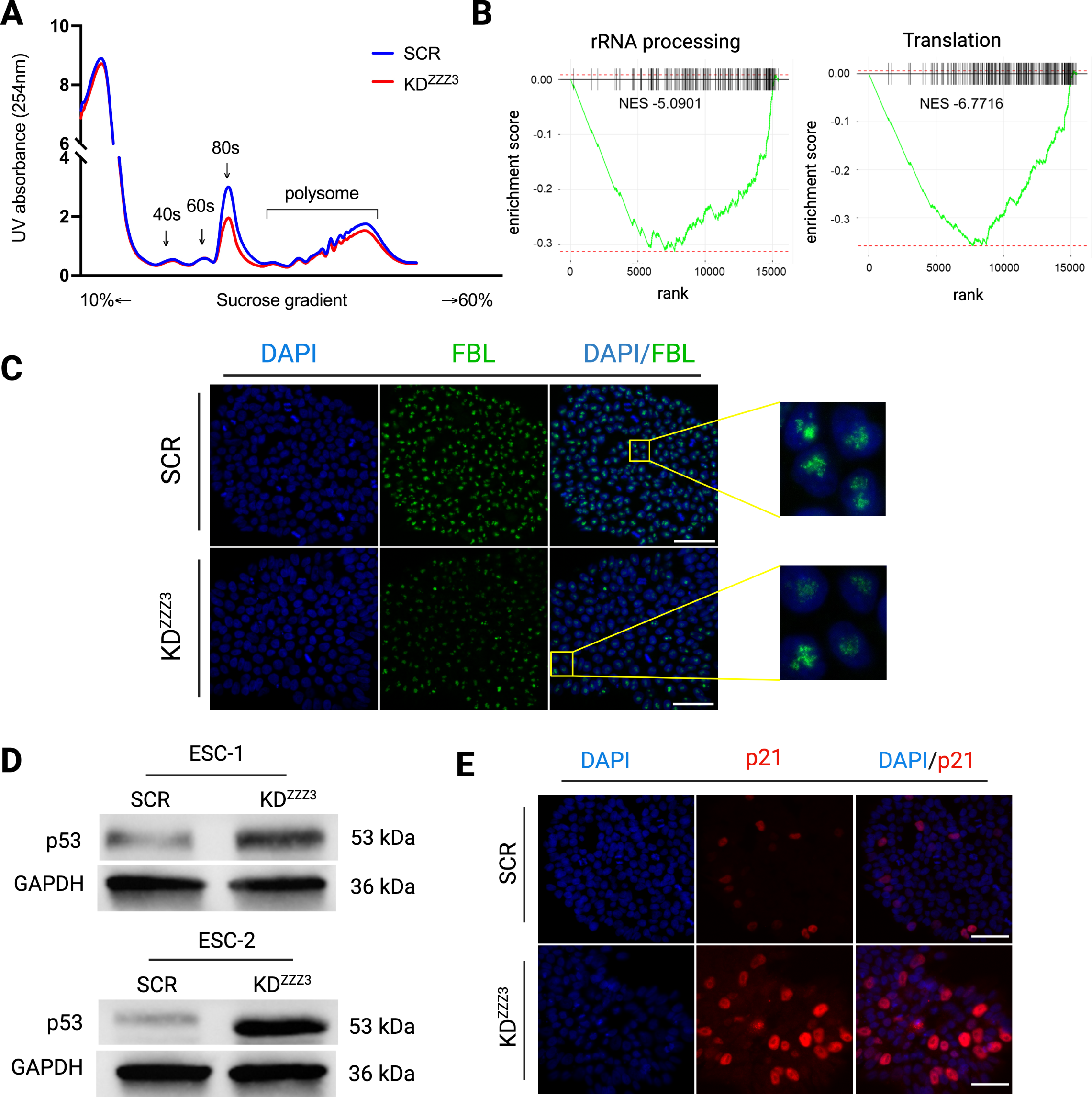
ZZZ3 knockdown leads to impairment in ribosome biogenesis and translation, and triggers nucleolar stress and p53 activation. **A** Polysome profiling absorbance, measured at 254 nm, of ZZZ3 KD cells and SCR control cell extracts fractionated on a 10-60% sucrose gradient. Peaks relative to free 40S and 60S subunits, 80S monosomes, and polysomes are indicated. **B.** GSEA enrichment plots relative to rRNA processing (NES = - 5.0901) and translation (NES = - 6.7716) gene sets in ZZZ3 KD *vs*. SCR ESCs. Adjusted *p*-value (*p*adj) is equal to 0.019 for both gene sets. **C.** Representative immunofluorescence images of the nucleolar fibrillarin (FBL) in ZZZ3 KD and control ESCs. Zoom-in images on the right highlight the reduced expression of FBL along with a more dispersed nucleolar structure in ZZZ3 KD ESCs. Nuclei were counterstained with DAPI. Scale bar, 50 µm. **D.** Quantitative immunoblot analysis of p53 showing an increased expression in ZZZ3 KD cells. GAPDH was used as loading control. Full-length blots are shown in Supplementary File S1. **E**. Representative immunofluorescence for p21 (red), a downstream target of p53, showing an increased expression in ZZZ3 KD ESCs.

### ZZZ3 controls cell proliferation and translation through that PI3K-AKT-mTOR axis

In the attempt to molecularly connect the reduced cell proliferation and impaired ribosome biogenesis observed in ZZZ3 KD ESCs, we focused on the ERK (Extracellular signal-regulated Kinase), and mammalian target of rapamycin (mTOR), the latter crucially involved in the regulation of translation initiation in mammalian cells. The PI3K-AKT-mTORC1 signaling pathways were significantly enriched by genes downregulated upon ZZZ3 depletion (Fig. 6A). The convergence of signals from the mTOR and PI3K pathways provides an integrated mechanism for cells controlling the rate of protein synthesis and cell growth (33). ERK is a member of the mitogen-activated protein kinase (MAPK) family, which plays a central role in cell proliferation. The phosphorylated ERK (p-ERK) serves as an indicator of the activation status of the ERK pathway and positively correlates with the rate of cell proliferation (34). We performed immunoblot analysis for total ERK1/2 and p-ERK (Thr202/Tyr204) and found that the expression of both is significantly reduced in ESCs with ZZZ3 KD (Fig. 6B). The mTOR pathway plays an important role in regulating various cellular processes, including ribosome biogenesis, by modulating processing of rRNA and the translation of ribosomal proteins, as well as activating the RNA polymerase I (Pol I) transcription machinery (35). mTOR activation stimulates protein synthesis primarily through the phosphorylation and regulation of downstream targets, including components of the translation machinery, such as ribosomal protein S6 kinase (S6K1 or p70S6 kinase) (36), leading to the phosphorylation of its downstream targets, such as the ribosomal protein S6 (RPS6), a critical step in the initiation of protein translation (37,38). Immunoblot analysis of the active phosphorylated Akt (Ser473), which activates mTOR by phosphorylation on its Ser2448, confirmed a reduced expression of both in ZZZ3 KD ESCs (Fig. 6C and 6D). The main downstream targets of mTOR signaling, phosphorylated P70S6K (T421/424) and phosphorylated RPS6 (Ser235) were also significantly reduced in ESCs with ZZZ3 depletion (Fig. 6E and 6F). The inactivation by phosphorylation of 4E-BP1 (eukaryotic initiation factor 4E-binding protein 1) on T37/42 is another key mechanism by which mTOR promotes protein synthesis (39). We found that the inactive form of 4E-BP1 is less expressed in ZZZ3 KD cells (Fig. 6G) as well as its downstream target eIF4E (eukaryotic initiation factor 4E) (Fig. 6H). S6K phosphorylates several targets involved in translation initiation, leading to increased ribosome biogenesis and translation initiation complex assembly. Altogether, these results suggest the PI3K, ERK, AKT and mTOR signaling pathways are less active in ESCs with ZZZ3 KD, thus providing a molecular link between ZZZ3 and reduced proliferation, ribosome biogenesis, and translation.

**Figure 6.**
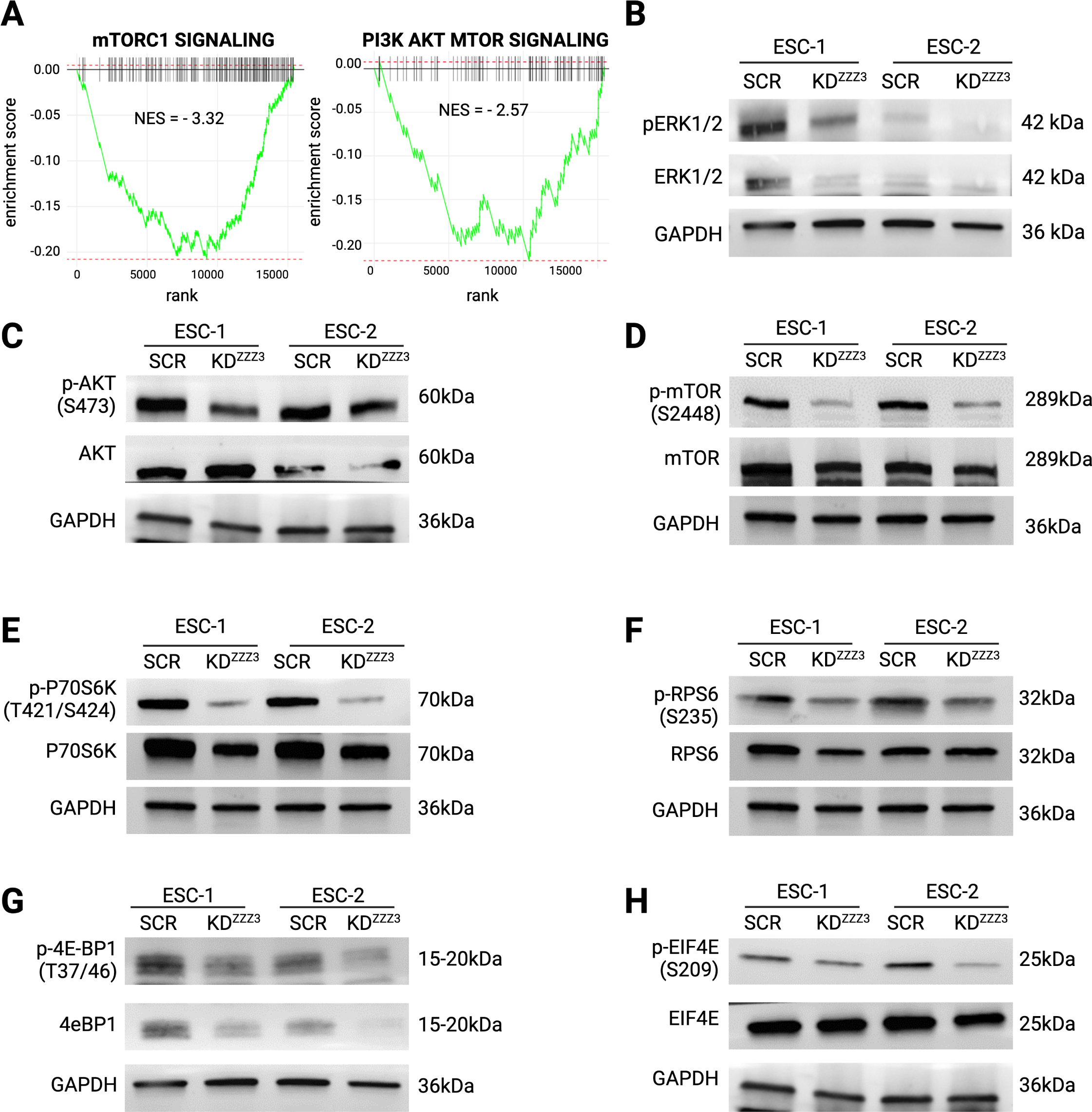
Knockdown of ZZZ3 reduces the PI3K-AKT-mTOR axis. **A.** GSEA enrichment plots relative to mTORC1 (NES = - 3.32) and PI3K/Akt/mTOR (NES = - 2.57) gene sets in ZZZ3 KD *vs*. SCR ESCs. Adjusted *p*-value (*p*adj) is equal to 0.0057 for both gene sets. **B-H**. Quantitative immunoblot analysis for phosphorylated ERK 1/2 (Thr202/Tyr204) and total ERK 1/2 (**B**); phosphorylated Akt (Ser473) and total AKT (**C**); phosphorylated mTOR (Ser2448) and total mTOR (**D**); phosphorylated P70S6 Kinase (Thr421/Ser424) and total P70S6 Kinase (**E**); phosphorylated S6 Ribosomal Protein (Ser235) and total RPS6 (**F**); phosphorylated 4E-BP1 (Thr37/46) and total 4E-BP1 (**G**); phosphorylated eIF4E (Ser209) and total eIF4E (**H**). GAPDH levels were assayed to confirm equal loading control in all immunoblot analysis. Full-length blots are shown in Supplementary File S1.

## Discussion

Pluripotent stem cells (PSCs), including embryonic stem cells (ESCs) and induced pluripotent stem cells (iPSCs), have the unique capability to self-renew while remaining competent to differentiate into any of the three germ layers (1,2,40). These features make human pluripotent stem cells extremely attractive for various biomedical applications including the understanding of early human developmental processes (41), disease modeling (42–44), drug discovery (45), regenerative medicine (46), and personalized medicine (47). For these applications to serve for basic and translational research, a thorough and comprehensive understanding of the molecular mechanisms controlling pluripotency is required. A large body of literature has mainly focused on transcriptional regulation of pluripotency (6–8) and mechanisms governing differentiation and cellular fate (48). However, PSCs as well as the embryo are highly proliferative. What are the mechanisms governing human PSCs proliferation? Must the two processes, proliferation and pluripotency, necessarily be coupled, or can they be independent? Studies conducted on diapaused mouse blastocysts and mouse ESCs suggest that pluripotency and proliferation can indeed be independent meaning that it is possible for mouse ESCs to maintain their pluripotent state without actively proliferating (10,12,13,49). mTOR was associated with high translational output in growing cells by direct phosphorylation and inactivating 4EBP1, which represses translation. Low levels of phospho-4EBP1 and phospho-Akt, a positive regulator of proliferation and metabolism, were found in paused and diapaused blastocysts (50). Altogether, these reports suggest that a significant reduction of protein synthesis takes place in diapaused blastocysts (49,51). Regulated translation comes into play as a mechanism through which stem cells can rapidly respond to cues from their microenvironment and adjust their protein synthesis patterns accordingly (52,53). The downregulation of translation is a key molecular mechanism underlying the dormancy of hPSCs; when hPSCs are in a dormant or paused state, they reduce their overall protein synthesis rate as a way to conserve energy and resources. Recently, it was reported that inhibition of mTOR can delay the progression of human blastoids and PSCs (54). Here, we focused on ZZZ3, whose function was associated with the regulation of genes encoding ribosomal proteins (RPGs) in the context of adenocarcinoma cell lines and mouse ESCs (14,15). We used a combination of proteomic, transcriptomic, and cellular analyses to investigate the role of ZZZ3 in human ESCs. To understand the specific function of ZZZ3 in human ESCs, we knocked down its expression and compared the characteristics of cells with reduced ZZZ3 expression to those with normal ZZZ3 expression. From a morphological point of view, ZZZ3 KD ESCs were similar to control cells and showed similar AP activity. The expression of pluripotency-associated markers such as *OCT4, SOX2, NONOG, MYC, LIN28A, LIN28B, BCOR, FOXO1, DPPA2, DPPA4, UTF1, KLF7* was also comparable. ZZZ3 depleted ESCs were able to form EBs and differentiate into the three germ layers in a fashion similar to SCR cells, suggesting that ZZZ3 deficiency does not impair pluripotency and differentiation capabilities of hESCs. On the other hand, ESCs are not only characterized by their pluripotency and differentiation capability but they are also highly proliferative. When looking at the growing rate, we noticed a substantial difference between ZZZ3 KD and SCR ESCs, as modified cells were significantly less proliferative than control cells and the expression of Ki67 was dramatically reduced in hESCs with ZZZ3 KD. Noteworthy, the ERK/PI3K/Akt/mTOR signaling, known to stimulate cell cycle progression, DNA replication, and cell growth, was significantly enriched by downregulated genes in ZZZ3 KD ESCs. Recent studies have begun to delineate the contribution of PI3K/AKT/mTOR signaling in preserving the ability of PSCs to self-renew and differentiate (55–58), and activation of the Akt pathway by phosphorylation on threonine 308 and serine 473 residues is known to promote cell survival and proliferation (59,60). Collectively, these results suggest that ZZZ3 is required for proper ESCs proliferation. Cell proliferation, growth, and protein synthesis are functionally and regulationally connected processes as proliferation cannot take place without proper increase in cell mass which, in turn, is dependent on proper protein synthesis (61). Interactome analysis in wild-type ESCs and transcriptome analysis in ZZZ3 KD and control ESCs allowed us to identify ribosome biogenesis, rRNA processing and metabolism, RNP complex biogenesis and assembly, RNA splicing, translation as overrepresented biological processes associated with ZZZ3 function. Here, we used polysome profile analysis to measure the translational activity in cells with ZZZ3 KD. Interestingly, we found a reduction of free ribosomes (80S) and polysome fractions, suggesting that modified cells have a lower translation efficiency, likely due to an impairment of ribosome biogenesis (5), which is a well-recognized canonical hallmark of cell proliferation and growth. The mTOR pathway is a key regulator hub that stands upstream of protein synthesis and plays a central role in coordinating various cellular processes, including cell growth, proliferation, metabolism, and protein synthesis. Activated mTOR phosphorylates and activates downstream targets that directly regulate protein synthesis. One of the most important targets is the ribosomal protein S6 kinase (S6K1 or p70S6 kinase) (62–64), which phosphorylates ribosomal protein S6 (RPS6) that promotes translation initiation and elongation (37). Immunoblot analysis of Akt, an upstream regulator and activator of mTOR, mTOR, and its targets RPS6 and S6K1, were significantly reduced in ZZZ3 depleted ESCs. The decreased levels of Akt, mTOR, and their targets in ZZZ3-depleted cells may explain the observed impairment in ribosome biogenesis and translation. Since mTOR is a key regulator of these processes, reduced mTOR activity could lead to disruptions in ribosome assembly and protein synthesis (64). ZZZ3 is mainly localized in the nucleoplasm and nucleoli, the main site of ribosome biogenesis, which represents a key point in cell growth, tightly linked to cell proliferation (65). Immunostaining for FBL, which is enriched in the dense fibrillar component (DFC) of the nucleolus, indicated that in ZZZ3 KD cells, the nucleolar structure is more dispersed compared to the nucleolar structure in SCR ESCs. This could imply that ZZZ3 plays a role in maintaining the organization or integrity of the nucleolus and could potentially be involved in processes related to ribosome biogenesis. Impaired ribosome biogenesis, in turn, can trigger nucleolar stress via a p53-dependent mechanism (32). In addition to its primary function in ribosome production, the nucleolus also acts as a regulatory hub for stress signals, and one of the key stress response pathways it influences is the activation of the p53 checkpoint (66). Active p53 exerts its protective mechanism by preventing cells from proliferating with damaged or incomplete ribosomes (67). In ZZZ3 KD cells both p53 and its downstream target p21 were dramatically increased, suggesting that nucleolar stress may occur upon ZZZ3 silencing. Overall, the main findings of our research can be summarized as follows: *1)* proliferation appears to be independent of pluripotency in hESCs *in vitro*; ZZZ3-KD hESCs adapted to a less proliferative, biosynthetic quiescent pluripotent state resembling mESCs diapause; *2)* ZZZ3 is crucial to maintain a high proliferative state of hESCs; it does so by ensuring proper ribosome biogenesis and translation likely through its interaction with the PI3K/Akt/mTOR pathway; *3)* ZZZ3 depletion correlates with an impairment of ribosome biogenesis and nucleolar integrity leading to nucleolar stress and activation of p53. Active p53 prevents the propagation of cells with compromised or unbalanced protein synthesis.

## Methods

### Cell culture

The human ES cell-lines (WA17 and RUES2; WiCell Research Institute, Madison, WI) were maintained in mTeSR1 Plus medium (STEMCELL Technologies, Vancouver, Canada) on plates coated with Matrigel (BD Biosciences, San Diego, CA) in a humidified incubator at 37°C and 5% CO_2_. Medium was replaced every two days and cells were split every 5 to 6 days using Gentle Dissociation reagent (STEMCELL Technologies, Vancouver, Canada). hES cells were routinely tested for Mycoplasma using the Mycoplasma PCR Detection kit (Applied Biological Materials, Richmond, Canada).

### Generation of ZZZ3 knockdown ES cells

hES cells with stable knockdown of ZZZ3 gene were generated by transfecting cells with PiggyBac (PB) transposon plasmids system with PB transposase expression vector pBase (kindly provided by Graziano Martello’s laboratory, University of Padua). Three PB U6-promoter driven shRNAs against ZZZ3 mRNA or scramble control were purchased from Vector Builder. For DNA transfection, hES cells were dissociated as single cells using TrypLE^TM^ Select Enzyme (1X) (ThermoFisher Scientific) and 250,000 cells were co-transfected with PB constructs (550 ng) and pBase plasmid (550 ng) using FuGENE HD transfection reagent (3.9 μl) (Promega), following the manufacturer’s instruction. Cells were cultured onto Matrigel-coated 12-well plate in mTeSR1 Plus medium (STEMCELL Technologies, Vancouver, Canada) with 10 μM Y27632 (ROCKi, Rho-associated kinase (ROCK) inhibitor, Selleckchem). After 48 h, transfected cells were selected with Hygromycin B (200 μg/ml, Thermo Fisher Scientific) diluted in mTeSR1 Plus for two weeks before performing experiments.

### Cell proliferation assay

hES cells were seeded on Matrigel-coated 12-well plates (3×10^4^ cells/well) in mTeSR1 Plus medium for the indicated time periods (24h, 48h, 72h, and 96h). Cells were dissociated with Accutase (Thermo Fisher Scientific) and counted with a cell counting chamber. Each experiment was performed independently three times.

### MTT assay

5,000 hES cells per well were seeded in triplicate onto a 96-well plate matrigel-coated. At 80% confluence, cells were treated with 0.5 mg/ml of 3-[4,5-dimethylthiazol-2-yl]-2,5-diphenyl-tetrazolium bromide (MTT) for 2 h. MTT solution was replaced with 2-propanol (Sigma-Aldrich) under agitation for 10 minutes. Absorbance was measured at 570 nm using a Varioskan LUX plate reader (Thermo Fisher Scientific).

### EBs formation assay

shNT and ZZZ3 knockdown ESCs were dissociated into single cells using StemPro Accutase (Thermo Fisher Scientific) and plated (200,000 cells/well) on low attachment 12-well plates coated with poly-2-hydroxyethyl methacrylate in mTeSR1 Plus (STEMCELL Technologies, Vancouver, Canada) medium with 10 μM Y27632 (Selleckchem) for three days. On day three, the medium was refreshed (without Y27632) every other day until day seven. Brightfield images of EBs were captured using an imaging system (DMi8, Leica Microsystems) at 5X magnification. EBs diameters were manually assigned using ImageJ software. The data set was then exported to Microsoft Excel for data analysis. EBs differentiation was performed as previously described in (16). Images of differentiated EBs (d28) were acquired at 63X magnification with imaging systems (DMi8), filter cubes and software from Leica Microsystems.

### Immunofluorescence and stainings

Immunofluorescence analysis was performed on Matrigel-coated glass coverslip in wells. Cells were fixed in 3.7% (vol/vol) formaldehyde for 15 minutes at room temperature, washed in PBS, permeabilized for 1 h at RT in PBS + 0.3% Triton X-100 (Sigma-Aldrich) (PBST), and blocked for 1 h at RT in PBST containing 10% of fetal bovine serum (FBS) (Thermo Fisher Scientific). Cells were subjected to immunostaining overnight at 4°C with primary antibodies (Table S3) diluted in blocking solution. After washing with PBS, cells were incubated with Alexa Fluor 488- or 594-conjugated secondary antibodies (Alexa, Life Technologies) (Table S3) for 1 h at RT. Nuclei were stained with DAPI (4′,6-diamidino-2-phenylindole) (Thermo Fisher Scientific). Finally, cells were mounted with DAKO Fluorescent Mounting Medium (Agilent), and images were acquired using Leica microscopy systems (DMi8 and Thunder DMi8) and Leica LAS X software (v.3.7.4.23463). All immunostainings analyses were performed using ImageJ software.

### Western blot analysis

Cells were washed with cold PBS and then scraped from the plate using RIPA buffer (150mM Sodium Chloride, 1% Triton x-100, 0.5% sodium deoxycholate, 0.1% SDS (sodium dodecyl sulfate), 50mM Tris hydrochloride, pH 8.0), supplemented with Halt™ Protease Inhibitor and Halt™ Phosphatase Inhibitor Cocktails (Thermo Fisher Scientific). The protein content was determined using Bradford (Bio-Rad) protein assay. Equal amounts of proteins (30-50 μg) were denatured in Laemmli sample buffer at 95°C for 5 minutes, separated on 4-20% Mini-PROTEAN TGX precast gels (Bio-Rad), and transferred onto nitrocellulose membranes (Bio-Rad) using a Trans-Blot® Turbo™ Transfer System (Bio-Rad). Blots were blocked in 5% non-fat milk for 1 h at room temperature and subsequently subjected to overnight primary antibody incubation at 4°C, followed by two quick rinses and three washes for 5 min in PBST (PBS + 0.1% Tween-20). Secondary antibody incubation was performed for 1 h at RT. Clarity™ Western ECL Blotting Substrates (Bio-Rad) was used to detect the HRP signal and the western blot images were collected using the Alliance™ Q9-Atom (Uvitec). Uncropped western blots images are shown in Supplementary figure S6. The information for specific antibodies is listed in Table S3.

### RNA extraction, reverse transcription, and quantitative real-time PCR

Total RNA was extracted using TRIzol Reagent (Thermo Fisher Scientific) following manufacturer’s instructions. Reverse transcription was performed using High-Capacity cDNA Reverse Transcription Kit (Thermo Fisher Scientific). Quantitative PCR analyses were performed in real time using a QuantStudio™ 7 Pro Real Time PCR system (Applied Biosystem) and SensiFAST SYBR Hi-ROX kit (Meridian Bioscience). Gene expressions were calculated following normalization to *GAPDH* (Glyceraldehyde 3-phosphate dehydrogenase) levels using the comparative Ct (cycle threshold) method. Statistical differences were calculated using two-tailed *t*-test or multiple unpaired *t*-test with Welch correction, with a significance of * *p* ≤ 0.05, ** *p* ≤ 0.01, and *** *p* ≤ 0.001. Data are presented as mean ± SEM from three independent experiments. The sequence of primers pair used in qRT-PCR analysis are the following: *OCT4 (*Fw-GGAGGAAGCTGACAACAATGAA; Rev-GGCCTGCACGAGGGTTT); *NANOG* (Fw-TGCAAGAACTCTCCAACATCCT; Rev-ATTGCTATTCTTCGGCCAGTT); *SOX2* (Fw-ATGCACCGCTACGACGTGA; Rev-CTTTTGCACCCCTCCCATTT); GAPDH (Fw-TCCTCTGACTTCAACAGCGA; Rev-GGGTCTTACTCCTTGGAGGC).

### RNA-Sequencing library preparation and analysis

Total RNA was extracted using TRIzol Reagents (Invitrogen, 15596026) according to the manufacturer’s protocol. For library preparation, the quantity and quality of the starting RNA were checked by Qubit and Bioanalyzer (Agilent). 1 μg of total RNA was subjected to poly(A) enrichment and library preparation using the TruSeq® Stranded mRNA Library Prep Kit (Illumina) following the manufacturer’s instructions. Libraries were sequenced on Illumina NextSeq 1000 System (paired-end 60+60 bp reads). After quality controls with FastQC (https://www.bioinformatics.babraham.ac.uk/projects/fastqc/), raw reads were aligned to the human reference genome (hg38/GRCh38) using STAR 2.7.1a (17) (with parameters– outFilterMismatchNmax 999 –outFilterMismatchNoverLmax 0.04). Gene expression levels were quantified with featureCounts v1.6.3 (18) (options: -t exon -g gene_name) using GENCODE 32 annotation. ZZZ3 is annotated in GENCODE as AC118549.1. Multi-mapped reads were excluded from quantification. Gene expression counts were next analyzed using the edge*R* package (19). After low expressed genes filtration (1 count per million (CPM) in less than 3 samples), normalization factors were calculated using the trimmed-mean of M-values (TMM) method implemented in the calcNormFactors function and CPM were obtained using normalized library sizes. Differential expression analysis was carried out by fitting a GLM to all groups and performing LF test for the pairwise contrasts considering the different ESC line in the formula ‘0 ∼ genotype + cell_line’. Genes were marked as significantly differentially expressed when having | logFC | > 1 and adjusted *p*-value < 0.05 (Benjamini-Hochberg *p*-value correction). Principal component analysis (PCA) of the expression dataset was performed using the Prcomp function implemented in the R stats package. Gene expression heatmaps with hierarchical clustering based on euclidean distance were generated using the ComplexHeatmap R package scaling logCPM values as Z-scores across samples (20). Gene set enrichment analysis was conducted by using GSEA software (21) on logFC*-log(p-value) ranked genes.

### Immunoprecipitation

Subconfluent cells were harvested and cross-linked using 0.05 mM DSP (Thermo Fisher Scientific) for 2 h on ice. The reaction was quenched using 20 mM Tris-HCl, pH 7.4 for 15 min on ice, followed by two washes with PBS. Cells were centrifuged at 5,000 rpm for 5 min at 4° C and resuspended in IP buffer (50 mM sodium phosphate pH 7.2, 250 mM NaCl, 0.1% Triton X-100, 0.1 nM ZnCl2) supplemented with protease and phosphatase inhibitor cocktails (Thermo Fisher Scientific). The cell suspension was then sonicated with a Diagenode Bioruptor (Settings: 10s ON, 10s OFF, high power) and centrifuged at 13,000 rpm for 20 min at 4° C. Protein concentration was determined using the Bradford assay. 1 mg of cell lysate was immunoprecipitated overnight at 4° C under rotation with Dynabeads Protein A (1.5 mg) (Thermo Fisher Scientific) and 10 µg anti-ZZZ3 antibody (Abcam) or 10 µg anti-IgG antibody (Diagenode). The next day, the immunoprecipitated complexes were washed three times with an IP buffer. Precipitated proteins were trypsin-digested for mass spectrometry analysis. For ZZZ3 interactome by nanoLC-MS/MS, immunoprecipitation using anti-ZZZ3 antibody was performed in biological quadruplicate.

### Mass spectrometry

ZZZ3 and its interactors were released from the magnetic beads by pre-digestion in 100 µl of digestion buffer (100 mM Tris-HCl pH 8.5, 0.1% Triton X-100, 400 ng trypsin Proteomics Grade (Sigma-Aldrich) for 15 min at 37°C with shaking (500 rpm). After on-bead pre-digestion by trypsin, the supernatant was collected and subjected to tryptic digestion. Proteins were reduced by 10 mM dithiothreitol (DTT) for 1 h at 37°C under shaking (500 rpm). Cysteines were alkylated by treatment with 24 mM iodoacetamide (IAA) for 1h at 37°C with agitation (500 rpm) in the dark. The alkylation reaction was quenched by adding 2 mM DTT (final concentration) for 30 min at 37°C. Finally, proteins were incubated with 200 ng trypsin overnight at 37°C with agitation to complete digestion (22). The resulting tryptic peptides (20 µl corresponding to approximately 20 µg of peptides) were purified by strong cation exchange (SCX) StageTips method (23) to remove eventual detergent residues. Before loading on the StageTips, the digested samples were acidified by adding 80% acetonitrile-0.5% formic acid (wash solution 2). The StageTip was conditioned by adding 20% acetonitrile-0.5% formic acid (wash solution 1, 50 µl) and 50 µl of W2. Peptides were loaded by slowly letting the fluid pass through one plug using a benchtop centrifuge. After 2 washes with 50 µl of Solution W2 and 50 µl of Solution W1 respectively, peptides were eluted by adding 7µL of 500 mM ammonium acetate-20% acetonitrile, diluted with 33 µl of formic acid 0.1% and analyzed by nanoLC-MS/MS.

### LC-MS/MS analysis

Peptides were separated by an Easy nLC-1000 chromatographic instrument coupled to a Q-Exactive “classic” mass spectrometer (both from Thermo Scientific, Bremen, Germany). All the LC-MS/MS analyses were carried using a linear gradient of 75 min at a flow rate of 230 nl/min on a 15 cm, 75 μm i.d., in-house-made column packed with 3 μm C18 silica particles (Dr. Maisch). The binary gradient was performed using mobile phase A (0.1% FA, 2% ACN) and mobile phase B (0.1% FA and 80% ACN). Peptide separation was obtained at a flow rate of 230 nL/min and ramped using from 3% B to 40% B in 60 min, from 40% to 100% in 13 min; the column was cleaned for 5 min with 100% of B. LC-MS/MS analysis were performed in data-dependent acquisition (DDA) using a top-12 method. Full scan m/z range was 350-1800, followed by MS/MS scans on the 12 most intense precursor ions. DDA analysis was performed with resolution for full MS scan of 70,000 and of 35,000 for MS/MS scan; the isolation window was 1.6 m/z. AGC target for full MS was 1e6 and of 1e5 for MS/MS scan. The maximum injection time was set to 50 ms for full MS scans and to 120 ms for MS/MS scans. HCD fragmentation at normalized collision energy of 25 and dynamic exclusion 20s.

### DDA data analysis

Raw files were processed in MaxQuant software (version 2.0.1.0) using the Andromeda search engine. The MS/MS spectra were searched against a human proteome database (downloaded in March 2016 and containing 42.013 sequences). For label free quantification (LFQ) analysis the following settings were used: Carbamidomethylation of cysteines as static modification, and oxidation of methionine and protein N-terminal acetylation as variable modifications. High confidence and unique peptides (minimum 1 peptide per protein group) were used for protein identification. Further parameters were set as follows: first and main search peptide tolerance, respectively 20 ppm and 4.5 ppm; isotope and centroid match tolerance, respectively 2 and 8 ppm; maximum number of missed cleavages, 2. Match between runs (MBR) option was activated, with match-time window set at 0.7 min and initial alignment window at 20 min. Only unique peptides were selected for quantification with a minimum ratio count of 1. For statistical analysis of MaxQuant output, the Perseus software (version 2.0.6.0) was used as follows: the LFQ intensity of proteins from the MaxQuant analysis were imported and contaminants, reverse identification, and proteins only identified by site were excluded from further data analysis. Data were transformed in logarithmic scale (log2). After filtering (at least three valid LFQ values in at least one group), remaining missing LFQ values were imputed from a normal distribution (width, 0.3; down shift, 1.8). Finally, for all the data sets, paired two-sample *t*-test was used to assess statistical significance of protein abundances using a 5% permutation-based FDR adjustment. A scaling factor was used for correction: s0: 0.2.

### Polysome profile

Before harvesting, cells were washed with ice-cold PBS supplemented with 100 μg/mL cycloheximide and resuspended in 1 mL lysis buffer (10 mM Tris-HCl pH7.4, 100 mM KCl, 10 mM MgCl2, 1% Triton-X 100, 1 mM DTT, 10 U/mL RNAseOUT (Invitrogen), 100 μg/mL of cycloheximide), and scraped. After 5 min of incubation on ice, cell lysate was centrifuged for 10 min at 14,000 rpm at 4°C. The supernatant was collected and protein content was determined by Bradford analysis (Bio-Rad). Equal protein amounts (4.2 mg) were loaded onto a 10-60% sucrose gradient obtained by adding 6 mL of 10% sucrose over a layer of 6 mL of 60% sucrose prepared in lysis buffer without Triton and containing 0.5 mM DTT, in a 12-mL tube (Polyallomer; Beckman Coulter). Gradients were prepared using a gradient maker (Gradient Master, Biocomp). Polysomes were separated by centrifugation at 37,000 rpm for 2.5 hours using a Beckman SW41 rotor. Twelve fractions of 920 µL were collected while polysomes were monitored by following the absorbance at 254 nm. Total protein was retrieved by 100% ethanol (EtOH) precipitation performed overnight, washed twice with 70% EtOH and analyzed by SDS-PAGE followed by Western blot.

### Statistical analysis

Experimental data are presented as means ± standard deviation of the mean (SEM) unless stated otherwise. Statistical significance was calculated unless stated otherwise by two-tailed unpaired *t-*test on two experimental conditions with p ≤ 0.05 considered statistically significant. Statistical significance levels are denoted as follows: *p ≤ 0.05; **p ≤ 0.01; ***p ≤ 0.001; ****p ≤ 0.0001. No statistical methods were used to predetermine sample size. Super exact test was performed to test the significance of the Venn diagram by the *R* package Exact.

## Data availability

The RNA-seq data from this study have been deposited on the GEO database.

## Acknowledgements

We would like to thank Prof. Graziano Martello (University of Padua, Italy) for his comments and for proofreading the draft. We would also like to thank Ms. Giorgia Lucia Benedetto and Ms. Desirèe Valente for their technical support.

## Author contributions

MLC, VL, EIP, and GC conceived and designed the experiments; MLC, VL, EIP, performed most of the experiments; SS, CZ, LS, DSM, and MSM performed the experiments; CC performed the bioinformatics analyses; RNA-Seq data analysis; MLC, VL, EIP, GC, and AP performed the data analysis and interpretation; EIP, VL, and GC drafted the manuscript; GC provided financial support; EIP and GC provided conceptual advice and supervised the study; GC and EIP wrote the final version of the paper. All authors read and approved the final manuscript.

---

**Figure S1.**
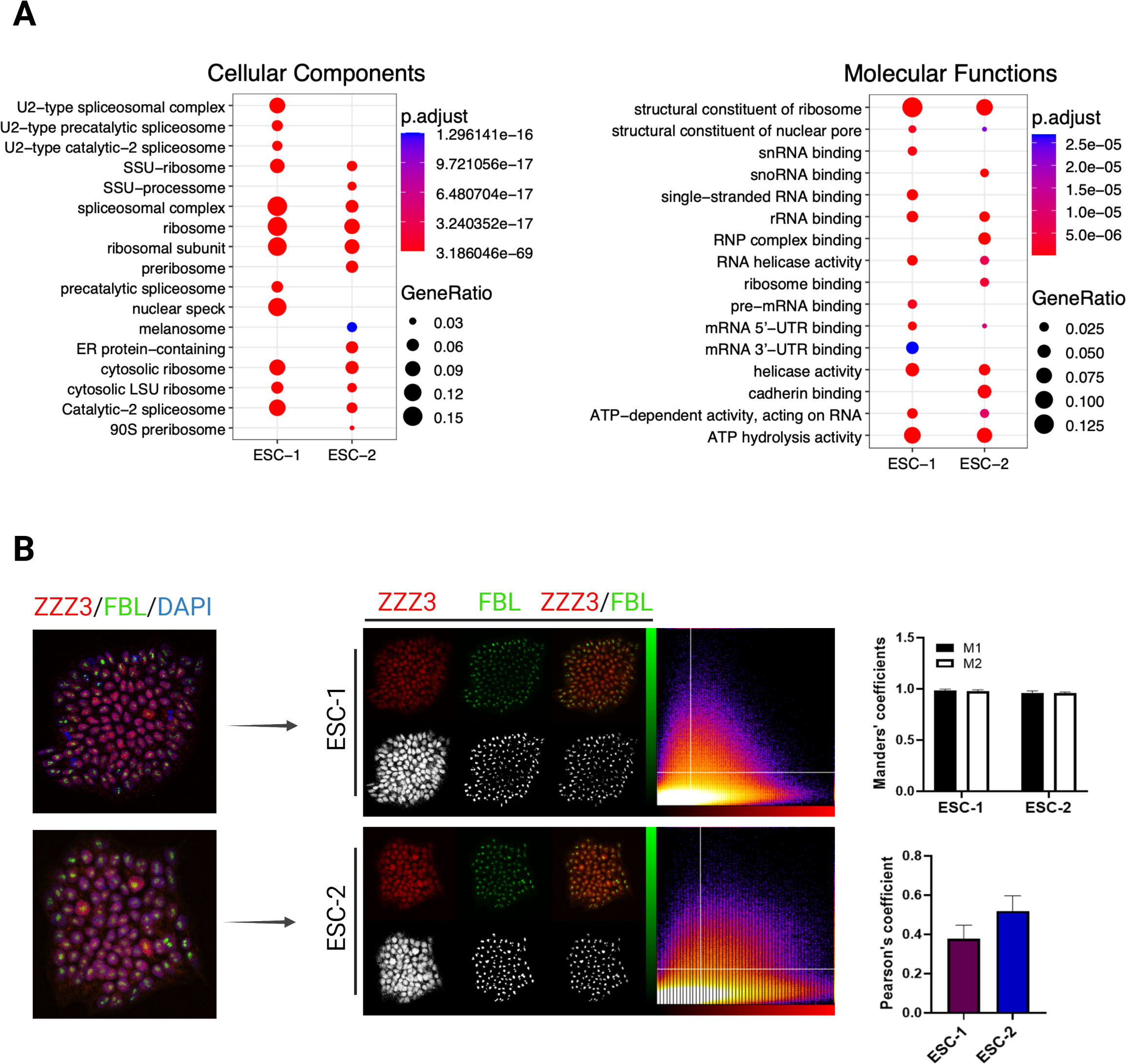
ZZZ3 is co-expressed with fibrillarin in the nucleolus and interacts with proteins involved in post-transcriptional processes and ribosome biogenesis. **A.** GO Cellular Component (left) and Molecular Function (right) for differentially expressed genes selected on the basis of the *p*-values (*p*-value < 0.05 corrected by using Benjamini-Hochberg procedure), and fold-change (FC ≥ 2.5). GO analysis was performed in *R* using the Bioconductor package. **B**. Immunofluorescence images of colocalization of ZZZ3 (red) and Fibrillarin (FBL, green) in wild-type ESC-1 and −2. The degree of colocalization was measured using the JACoP BIOP colocalization plugin of Image*J* software to calculate the Pearson correlation coefficient (PCC) and the Mander’s overlap coefficient (MOC) (M1-M2) on a total of 7 fluorescence images of both ESC lines analyzed. The intensity-based correlation analysis returned positive values of both PCC and MOC, indicating a strong co-localization of ZZZ3 and FBL in the nucleolus of wild-type ESCs.

**Figure S2.**
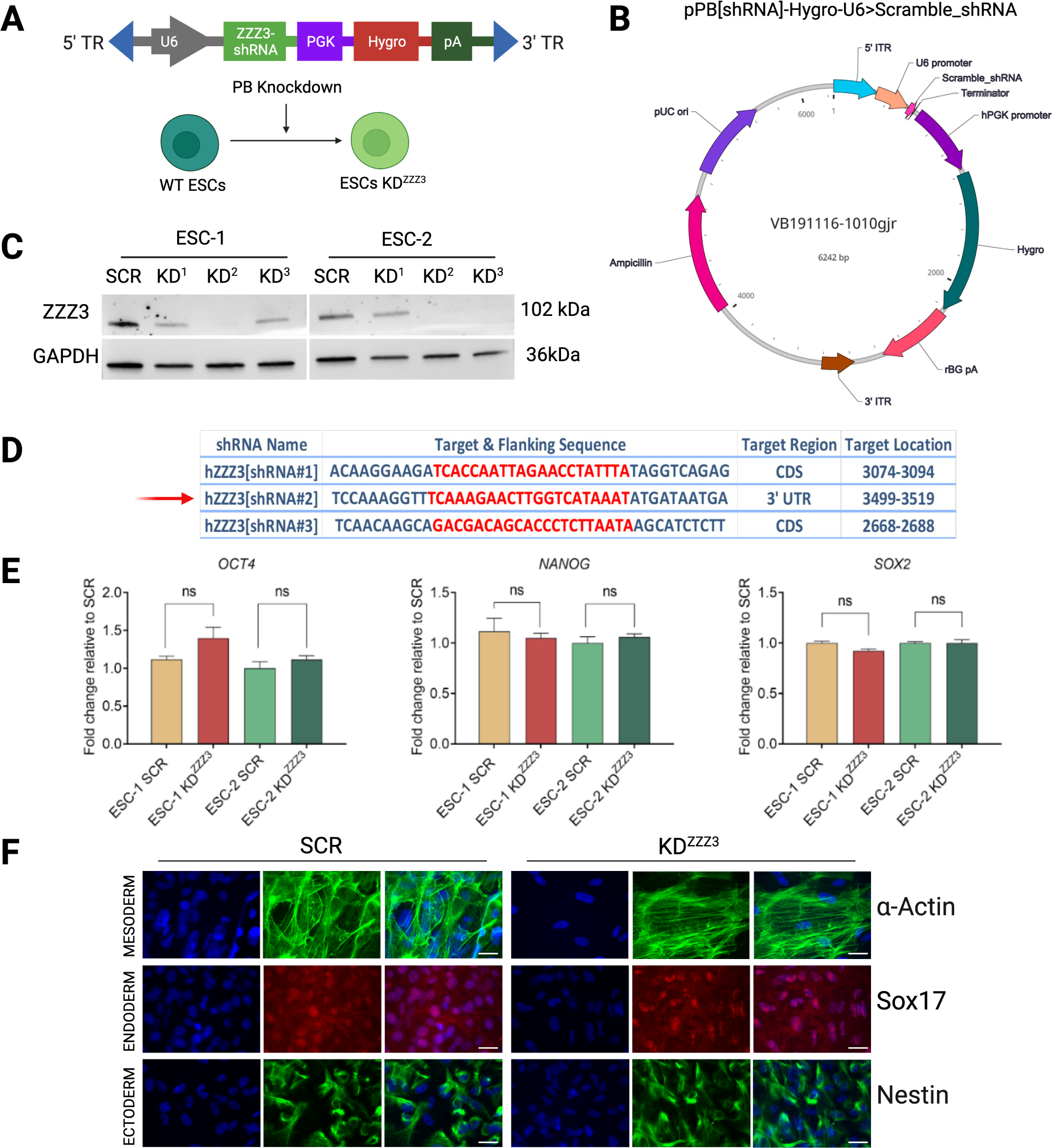
ZZZ3 knockdown was achieved by PiggyBac-mediated shRNA transfection and does not diminish the pluripotency of ESCs. **A.** Schematic representation of PiggyBac vector expressing shRNA against the human ZZZ3 gene under the control of U6 promoter used to generate stable ZZZ3 knockdown in hESCs. **B.** Map of the pPB[shRNA]-Hygro-U6>scramble_shRNA piggyBac vector. **C.** Immunoblot analysis was used to validate the ZZZ3 knockdown efficiency of three different shRNAs (KD^1^, KD^2^, KD^3^) in ESC-1 and ESC-2. ESCs transfected with shZZZ#2 (KD^2^) were used in this study. GAPDH was used as a loading control. Full-length blots are presented in Supplementary File S1. **D**. Table showing the sequence of the three shRNAs used to obtain hZZZ3 knockdown (hZZZ3[shRNA#1], hZZZ3[shRNA#2], hZZZ3[shRNA#3]). The target sequence, region and location of each shRNA on ZZZ3 gene sequence are indicated. **E.** Quantitative RT-PCR analysis of the core pluripotency factors *OCT4*, *NANOG*, *SOX2* (ns = not significant). Data are shown as mean ± SEM of three independent experiments and unpaired two-tailed *t*-test was calculated *vs.* SCR cells. **F.** Immunostaining of spontaneous differentiation into germ-layer specific cell types by means of EB formation. This representative figure of EBs at day 28 of differentiation shows the staining of the ZZZ3 KD ESCs and SCR cells for *α*-ACTIN (mesoderm-specific), SOX17 (endoderm-specific), and NESTIN (ectoderm-specific). Scale bar, 50 μm.

**Figure S3.**
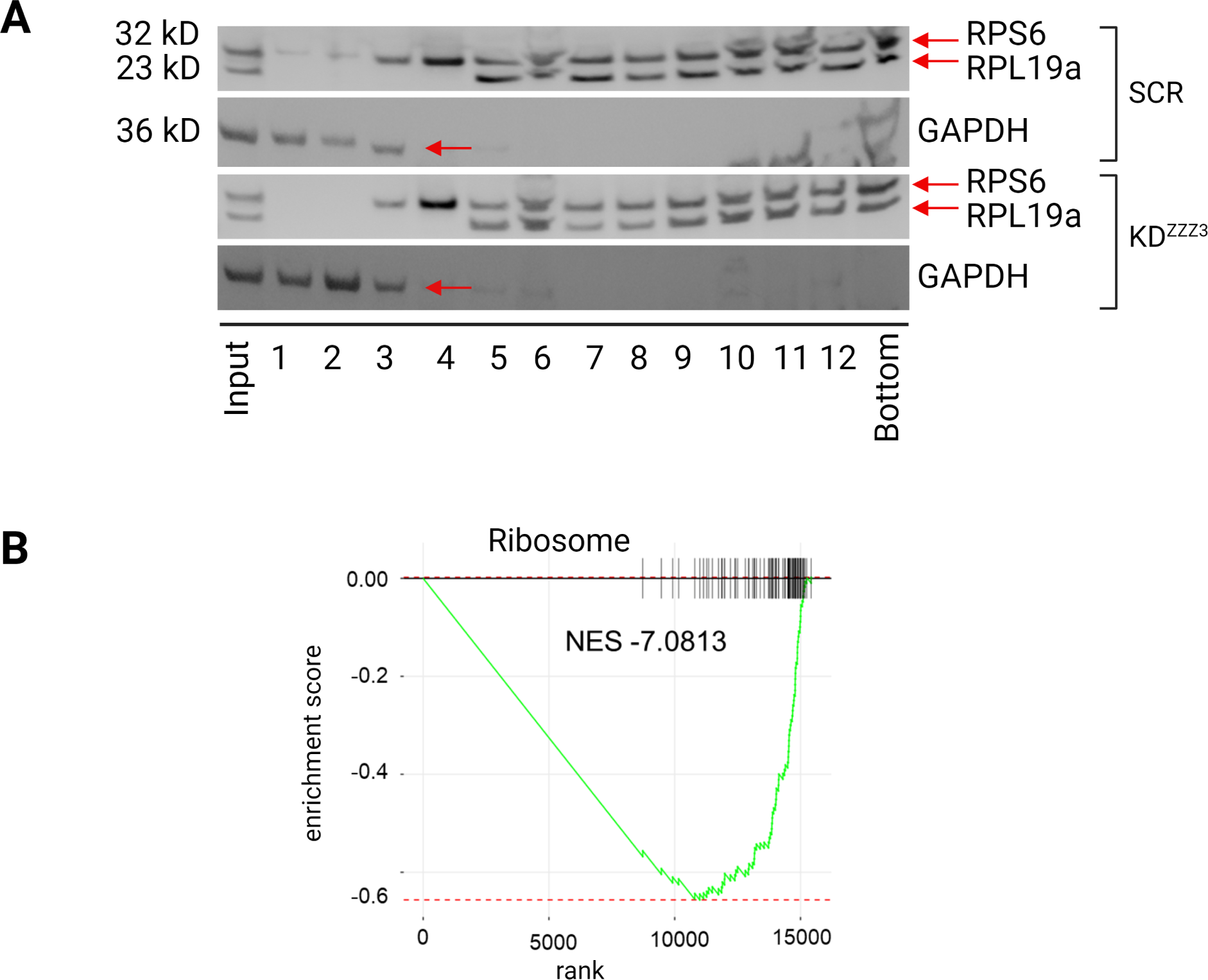
Many downregulated genes in ZZZ3 KD ESCs are enriched in the ribosome pathway. **A.** Immunoblot analysis of equal volumes of whole-cell polysomal fractions (see polysome profile in Fig. 5A). RPS6 and RPL19a are shown as references for small (40S) and large (60S) ribosomal subunits, respectively. GAPDH was used as a marker of proteins not associated with ribosomes. The numbers below images refer to different fractions: free ribosome (1-3), 40S subunit (4), 60S subunit (5), 80S monosomes (6), and polysomes (7-12). Full-length blots are presented in Supplementary File S1. **B**. GSEA enrichment plot (KEGG) relative to Ribosome pathway showing the significant decrease in the enrichment score (NES = - 7.081) for this gene set in ZZZ3 KD *vs*. SCR ESCs (*p*adj equal to 0.026).

**File S1.** Uncropped full-length gels.

**Table S1**. Gene Ontology (GO) interactomic data_ wild type ESC-1 and ESC-2.

**Table S2**. Differentially expressed genes (DEGs)_RNA-Seq ZZZ3 KD ESCs *vs*. SCR control ESCs.

**Table S3.** List of antibodies.

| ***Antibody*** | ***Host species*** | ***Dilution*** | ***Cat.No*** | ***Company*** |
| --- | --- | --- | --- | --- |
| ZZZ3 | Rabbit | 1:2000 | ab118800 | Abcam |
| Nanog | Rabbit | 1:1000 | PA1-097 | Invitrogen |
| Oct4 | Rabbit | 1:1000 | bs-1111R | Bioss Antibodies |
| pERK1/2 (Thr202/Tyr204) | Rabbit | 1:1000 | 9101 | Cell Signaling Technology |
| ERK1/2 | Mouse | 1:1000 | 9107 | Cell Signaling Technology |
| p53 | Mouse | 1:500 | sc-393031 | Santa Cruz Biotechnology |
| RPL19a | Mouse | 1:1000 | sc-100830 | Santa Cruz Biotechnology |
| Phospho-RPS6 (S235) | Rabbit | 1:1000 | ab227005 | Abcam |
| Ribosomal Protein S6 | Mouse | 1:2000 | sc-74459 | Santa Cruz Biotechnology |
| Phospho-Akt (Ser473) (D9E) | Rabbit | 1:2000 | 4060S | Cell Signaling Technology |
| Akt | Rabbit | 1:1000 | 9272S | Cell Signaling Technology |
| Phospho-mTOR (Ser2448) | Rabbit | 1:1000 | 5536S | Cell Signaling Technology |
| mTOR | Rabbit | 1:1000 | 2983S | Cell Signaling Technology |
| Phospho-4E-BP1 (Thr37/46) | Rabbit | 1:1000 | 2855 | Cell Signaling Technology |
| 4E-BP1 | Rabbit | 1:1000 | 9452 | Cell Signaling Technology |
| Phospho-eIF4E(Ser209) | Rabbit | 1:1000 | 9741 | Cell Signaling Technology |
| eIF4E | Rabbit | 1:1000 | 9742 | Cell Signaling Technology |
| Phospho-p70 S6 Kinase (Thr421/Ser424) | Rabbit | 1:1000 | 9204 | Cell Signaling Technology |
| p70 S6 Kinase | Rabbit | 1:1000 | 2708 | Cell Signaling Technology |
| Anti-acetyl-Histone H3 (Lys9) | Rabbit | 1:500 | S-06-942 | Sigma |
| GAPDH | Rabbit | 1:1000 | bs10900R | Bioss Antibodies |
| Actin | Goat | 1:500 | sc-1616 | Santa Cruz<br>Biotechnology |
| Peroxidase AffiniPure Donkey Anti-Rabbit IgG (H+L) |  | 1:10000 | 711-035-152 | Jackson Immuno<br>Reaserch |
| Peroxidase AffiniPure Sheep Anti-Mouse IgG (H+L) |  | 1:10000 | 515-035-062 | Jackson Immuno<br>Reaserch |
| Peroxidase AffiniPure Rabbit Anti-Goat IgG (H+L) |  | 1:5000 | 305-035-045 | Jackson Immuno<br>Reaserch |

List of antibodies for IF.
| ***Antibody*** | ***Host***<br>***species*** | ***Dilution*** | ***Cat.No*** | ***Company*** |
| --- | --- | --- | --- | --- |
| ZZZ3 | Rabbit | 1:100 | Pa5-84224 | Invitrogen |
| Fibrillarin | Mouse | 1:500 | ab4566 | Abcam |
| Nanog | Goat | 1:200 | AF1997 | R&D System |
| Oct4 | Mouse | 1:200 | 75463 | Cell Signaling<br>Technology |
| Nestin | Mouse | 1:1000 | 60091 | Stem Cell<br>Technology |
| Alpha-actin | Mouse | 1:500 | 14-9760-82 | Invitrogen |
| Sox17 | Goat | 1:20 | AF1924 | R&D System |
| Ki67 | Rabbit | 1:400 | 9129S | Cell Signaling<br>Technology |
| p21 | Rabbit | 1:400 | 2947S | Cell Signaling<br>Technology |
| Phospho-Histone H2A.X (Ser139) | Rabbit | 1:500 | 2577S | Cell Signaling<br>Technology |
| Anti-Rabbit IgG Alexa Fluor 594 |  | 1:500 | A-11012 | Invitrogen |
| Anti-Rabbit IgG Alexa Fluor 488 |  | 1:500 | A-11008 | Invitrogen |
| Anti-Goat IgG Alexa Fluor 594 |  | 1:500 | A-11058 | Invitrogen |
| Anti-Mouse IgG Alexa Fluor 488 |  | 1:2000 | A-11001 | Invitrogen |

